# Self-association features of NS1 proteins from different flaviviruses

**DOI:** 10.1101/2022.03.15.484429

**Authors:** Sergio A. Poveda-Cuevas, Catherine Etchebest, Fernando L. Barroso da Silva

## Abstract

Flaviviruses comprise a large group of arboviral species that are distributed in several countries of the tropics, neotropics, and some temperate zones. Since they can produce neurological pathologies or vascular damage, there has been intense research seeking better diagnosis and treatments for their infections in the last decades. The flavivirus NS1 protein is a relevant clinical target because it is involved in viral replication, immune evasion, and virulence. Being a key factor in endothelial and tissue-specific modulation, NS1 has been largely studied to understand the molecular mechanisms exploited by the virus to reprogram host cells. A central part of the viral maturation processes is the NS1 oligomerization because many stages rely on these protein-protein assemblies. In the present study, the self-associations of NS1 proteins from Zika, Dengue, and West Nile viruses are examined through constant-pH coarse-grained biophysical simulations. Free energies of interactions were estimated for different oligomeric states and pH conditions. Our results show that these proteins can form both dimers and tetramers under conditions near physiological pH even without the presence of lipids. Moreover, pH plays an important role mainly controlling the regimes where van der Waals interactions govern their association. Finally, despite the similarity at the sequence level, we found that each flavivirus has a well-characteristic protein-protein interaction profile. These specific features can provide new hints for the development of binders both for better diagnostic tools and the formulation of new therapeutic drugs.

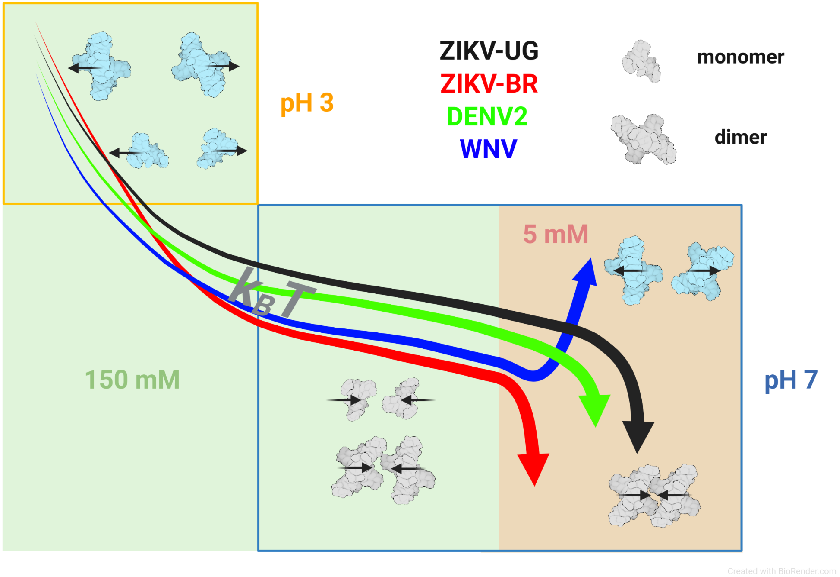

## 1. INTRODUCTION

The World Health Organization has reported in recent years viruses such as influenza H1N1, Chikungunya, Ebola, Lassa fever, MERS-CoV, Zika (ZIKV), Dengue (DENV), or West Nile (WNV) with epidemic or pandemic capacity (Trovato et al., 2020). The most recent outbreak by SARS-CoV-2 responsible for the Covid-19 pandemic, for example, has produced more than 452 million infections and more than 6 million deaths to date (World Health Organization, 2022). This ongoing crisis placed on high alert the health and economic worldwide systems, and reinforced the damage that these biological entities could have on a global scale. Through mutation processes, any virus can develop new variants resulting in discrepant pathological effects (Fields, 2014). In most cases, probably due to selective pressure, mutations can increase the viral mortality [as seen, for example, for Covid-19 or hemorrhagic dengue fever (DHF) by SARS-CoV-2 and DENV respectively (Gubler, 1998; Hu et al., 2021)] or can produce congenital damage [*e*.*g*., microcephaly by ZIKV (Mlakar et al., 2016)] The concern may be even greater considering the space-time coexistence of different viral species (Beck et al., 2013; Bhatt et al., 2013; Plourde and Bloch, 2016). For instance, in Brazil, up to 10 different species of flaviviruses have been reported in the last decades (Figueiredo, 2000). Many of these species affect the health of urban populations and there are no totally efficient serological tests due to the heterotypic immune responses associated with flaviviruses (Musso and Desprès, 2020).

The *Flavivirus* genus is composed of +ssRNA viruses with a genome of ∼11 kb that translates into a polyprotein. Through viral and host proteases the polyprotein is cleaved producing three structural proteins (prM, Env, and C) and seven non-structural (NS) proteins (NS1, NS2A, NS2B, NS3, NS4A, NS4B, NS5) (Neufeldt et al., 2018). The NS1 protein is involved in regulatory functions and different viral processes, such as replication, virulence, and pathogenicity. Consequently, NS1 is one of the most relevant clinical targets for flavivirus differentiation (Muller and Young, 2013; Watterson et al., 2016). It has also been suggested as a target for vaccine development due to the low risk of antibody-dependent enhancement (ADE) mechanism (Li et al., 2018; Sridhar et al., 2018; Wessel et al., 2021). Nevertheless, NS1 is well known to produce autoantibodies, which can promote inflammation and/or apoptosis into the host (Muller and Young, 2013). The first stage of NS1 in the endoplasmic reticulum (ER) shows a tendency to form dimeric structures with more liposoluble characteristics compared to its monomeric state (Winkler et al., 1989). The dimeric NS1 can interact with membranes (mNS1) (Noisakran et al., 2008), where it later participates as a cofactor in early events of viral RNA replication, virion assembly, and particle release (Scaturro et al., 2015). On the other hand, another portion of mNS1 is trafficked through the Golgi apparatus where it is secreted as a hexamer (sNS1) to be ready to interact with the host’s immune system (Flamand et al., 1999; Gutsche et al., 2011; Watterson et al., 2016). Yet, the NS1 dimer can also do so the same thing, showing that hexamerization is not a strict requirement for secretion (Watterson et al., 2016). Furthermore, the transition of a dimer to a hexamer implies the formation of tetrameric intermediates which are essential for the formation of sNS1 (Flamand et al., 1999). This viral toxin exhibits a high versatility that could be influencing the differential tropism among viral strains or species (*i*.*e*., at the intraspecific or interspecific level, respectively). Inflammatory processes during DHF by DENV, the overcoming of the blood-brain or placental barriers (by the neurotrophic WNV or ZIKV, respectively) imply the presence of sNS1 as an important step in immune activation or endothelial damage (Basu and Chaturvedi, 2008; Jurado et al., 2016; Neal, 2014; Puerta-Guardo et al., 2019; Watterson et al., 2016).

Some experimental and computational studies have focused on understanding the oligomerization processes of NS1 proteins as a possible therapeutic strategy. Site-directed mutagenesis assays have shown that there are an attenuation and virulence reduction for DENV, ZIKV, Kunjin, and Murray Valley encephalitis viruses when the NS1 dimerization is affected (Clark et al., 2007; Hall et al., 1999; Lin et al., 1998; Wang et al., 2017). Electrophoresis studies by Falconar and Young (1990) found that dimerization for DENV2 occurs for pH regimes greater than 3 and less than 7.4, while tetramerizations were reported at pH 8 and 8.5, for the NS1 of DENV2 and DENV1, respectively (Flamand et al., 1999; Muller et al., 2012). Nevertheless, no data is yet available to access the effects from their differences at the protein sequence level. Also, it is not clear if the presence of lipids is a fundamental need (Winkler et al., 1989). To deeply understand how flaviviruses work, it is essential to dissect this molecular mechanism for different species and strains.

Computer simulations are an interesting approach to unveil the phenomena that control the NS1 self-association. It allows testing specific steps at the molecular level under a well-controlled condition and without any risk of contamination and/or external interference typically at lower costs. Individual aspects of the interplay of several factors can be one by one tested. Molecular dynamics studies, for instance, have revealed that the β-hairpins of the β-roll domain are crucial for the interchain fit and stability of the NS1 dimers (Gonçalves et al., 2019; Menezes et al., 2020; Poveda-Cuevas et al., 2021). It has also been shown that the monomer-monomer association is guided by salt bridges or hydrogen bonds (Gonçalves et al., 2019), validating the importance of electrostatics in these biomolecular interactions. This also implies that pH is a thermodynamic variable that can trigger the association process (Poveda-Cuevas et al., 2020, 2018; Qiao and Olvera de la Cruz, 2020). Indeed, it is worth mentioning that the role of electrostatic interactions in complexation processes is not limited to flaviviruses. The phenomena produced by these interactions have been studied in other viral genders such as *Betacoronavirus* (Giron et al., 2021, 2020), *Chlorovirus* (Xian et al., 2019), *Bromovirus* (Suci et al., 2005), etc.

There is a differential electrostatic profile among flaviviruses despite their structural and sequence conservation (20%–40% identity and 60%–80% similarity) (Brown et al., 2016; Freire et al., 2017; Song et al., 2016; Xu et al., 2016). Interestingly, this also happens on Ugandan (UG) and Brazilian (BR) ZIKV variants from African and Asian origins, respectively. As previously shown via constant-pH (CpH) Monte Carlo (MC) simulations, the *pK*_*a*_s of ionizable groups from the NS1_ZIKV-UG_ suffer a displacement (or Δp*K*_*a*_) when compared with values from strains phylogenetically newer (*i*.*e*., NS1_ZIKV-BR_) (Poveda-Cuevas et al., 2018). In spite of their similar sequences, titratable residues are affected in a specific way for each NS1 protein at their three-dimensional configuration (Poveda-Cuevas et al., 2020, 2018). As it can be anticipated, such physical feature has consequences on three biological interface regions responsible for the *i*) oligomerization, *ii*) membrane binding, and *iii*) antibody interactions (Poveda-Cuevas et al., 2018).

The present work aims to study the self-association of four NS1 flaviviruses: a) NS1_ZIKV-UG_, b) NS1_ZIKV-BR_, c) NS1_DENV_ (specifically variant 2, NS1_DENV2_), and d) NS1_WNV_ at different pH conditions. Through CpH coarse-grained (CG) models solved via MC simulations, free energy derivatives are calculated in order to characterize/describe, dimerization and tetramerization processes of the NS1 protein. The hexamerization process is not studied here due to its multifactorial contributions: different types of biomolecules such as lipids and sugars could be necessary to stabilize the three dimers NS1, auxiliary proteins could manage the hexameric formation, or the hexamers are generated during a lipid droplet process nascent in the membrane of the Golgi apparatus (Gutsche et al., 2011; Somnuke et al., 2011). Thus, this would increase the complexity of the models far from the computational capacities. Further reasons for the choice of only these two simpler oligomeric states (*i*.*e*., dimers and tetramers) are deeply discussed in previous work (Poveda-Cuevas et al., 2018).

The present paper is organized as follows. In the next section, we describe the methodological aspects starting with the molecular models. Next, the numerical simulations used to solve the model and details of the different sets of calculations are described together with the main physical properties sampled. Subsequently, in the Results and Discussion section, the CG model is validated by comparing the theoretical outcomes with available electrophoresis and crystallography data. In the next step, the dimerization and tetramerization processes for the four NS1 proteins under relevant pHs and salt conditions were evaluated. Special attention was given to the self-association at the inter- and intraspecific levels. The last section aims to understand the physicochemical reasons that govern the NS1 oligomerization and the discrepancies among the flaviviruses. Lastly, how these factors could have a relationship with the biological aspects observed for each viral species is discussed.

## 2. METHODOLOGY

### 2.1. Structural modeling

Atomistic coordinates for the three-dimensional structures from NS1_ZIKV-UG_, NS1_ZIKV-BR_, NS1_DENV2_, and NS1_WNV_ deposited in the RCSB Protein Data Bank (PDB) (Berman et al., 2000) were used as input for the simulations. X-ray structures are available at the PDB and identified with ids 5K6K (1.89Å, pH 8.5) (Brown et al., 2016), 5GS6 (2.85Å, pH 8.5) (Xu et al., 2016), 4O6B (3.00Å, pH 5.5), and 4O6D (2.60Å, pH 5.5) (Akey et al., 2014) for NS1_ZIKV-UG_, NS1_ZIKV-BR_, NS1_DENV2_, and NS1_WNV_, respectively. The experimental resolution and pH conditions for each of these coordinates are the numbers given in parenthesis. The two NS1_ZIKV_ protein models employed here were selected based on our previous study (Poveda-Cuevas et al., 2018). Missing residues in the loop regions were completed through an appropriate homology modeling protocol (Poveda-Cuevas et al., 2021, 2018). The crystallographic structures of NS1_ZIKV-UG_ and NS1_ZIKV-BR_ are homodimers where one chain is fully resolved in the experiment and the other is not. Hence, the complete chain was used as a template to model the second chain with missing residues. The modeling was done through the “Modeller” program that is integrated into UCSF Chimera 1.11.2 (Pettersen et al., 2004). For each NS1_ZIKV_ five models were built and the structure with the best DOPE was selected (Shen and Sali, 2006). Afterwards, the PDB files were edited before the simulation by deleting all water molecules and heteroatoms.

The dimers NS1_DENV2_ (PDB id 4O6B) and NS1_WNV_ (PDB id 4O6D) also have missing residues in loop regions. The missing regions of the structure 4O6B (DENV2) are 8-10_chain_A_, 107-130_chain_B_, 108-128_chain_A_, 159-165_chain_A_, 162-163_chain_B_, and 350-352_chain_A/B_. Regarding the crystal given by the PDB id 4O6D (WNV), the missing parts are 108-128_chain_A_ and 108-128_chain_B_. Because both chains in these two structures are incomplete we can not follow the same strategy as implemented for the two NS1_ZIKV_. For this part of the study, a more advanced homology modeling protocol was applied. This process involved multiple templates (Larsson et al., 2008). Thereby, we start from the query sequence of the dimeric NS1_DENV2_ with the respective 704 amino acids. Later, three templates in the dimeric state were selected: *a)* the coordinates for NS1_DENV2_ given by the PDB id 4O6B (with missing residues but 100% of identity sequence when compared to the query sequence and excluding gaps), *b)* the NS1_ZIKV-UG_ (without any missing amino acids), and *c)* the NS1_ZIKV-BR_ (without any missing residues). The two structures of items *b* and *c* have some differences, especially in some loop regions (see Figure S1) (Poveda-Cuevas et al., 2021). Since both atomic arrangements of NS1_ZIKV-UG_ and NS1_ZIKV-BR_ are probable, they should be used simultaneously in order to increase the chance of obtaining structures for NS1_DENV2_ with more favorable energies (Fernandez-Fuentes et al., 2007; Larsson et al., 2008). As a first stage in the homology modeling, the protein sequences of the three templates were extracted to make alignments with the query sequence. For this process, the python scripts “salign.py” and “align2d_mult.py” (available in https://salilab.org/modeller/) were employed. From the multiple alignment sequence, ten 3D structure models with complete parts were built with the python script “model_mult.py” (also available in the previous web URL). The 3D structural model with the best DOPE score was used for our simulations (Shen and Sali, 2006). Regarding NS1_WNV_, the same strategy was followed as for NS1_DENV2_. However, the structure of item *a)* was replaced by the uncompleted crystal coordinates given by PDB id 4O6D. All scripts were executed with Modeller 9.24 (Webb and Sali, 2014) and default parameters.

The molecular representation of all NS1 proteins after all modeling steps above described is given in Figure 1. Note the differences in the location and spatial distribution of the titratable residues for each system. Monomers were prepared by the simple deletion of one given chain. Chain A was selected for most models except for NS1_ZIKV-UG_ where chain B was chosen. All final atomistic coordinates are available for download as Supplementary Materials (https://zenodo.org/record/5608521). The monomeric and dimeric states are referenced in this work with the subscripts “m” and “d”, respectively. After this initial preparation, the atomic coordinates were converted into the simplified CG molecular model as described in the next subsection.

**Figure 1.**
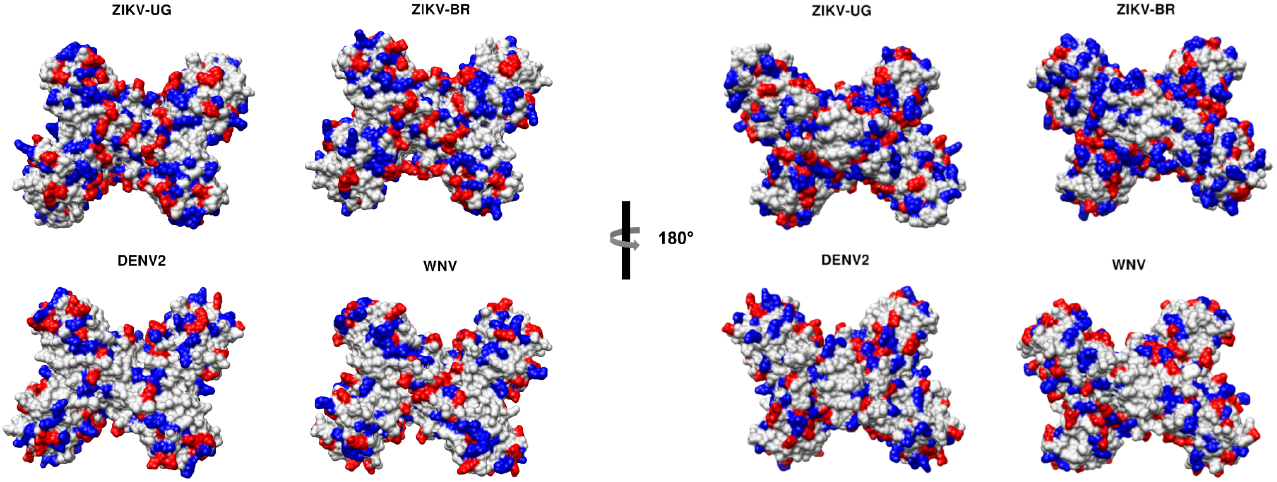
Molecular representation of the NS1 proteins (dimers) for the flaviviruses ZIKV-UG (PDB id 5K6K), ZIKV-BR (PDB id 5GS6), DENV2 (PDB id 4O6B), and WNV (PDB id 4O6D) in a surface representation. Positive and negatively charged amino acids are represented in blue and red, respectively.

### 2.2. Structural analysis

Additional calculations were done in order to compare structural features of the four NS1_flavivirus_: *a)* all possible pairs of the whole dimer structures were overlapped and computations of the root mean square deviation (RMSD) using the alpha carbons (C_α_) were performed, *b)* from the overlapped dimers of item *a*, the RMSDs per residue were estimated, *c)* the percentage of identity among the NS1 protein sequences were also calculated, and *d)* volumes of the NS1 proteins in the monomeric and dimeric states were estimated. The items *a* to *d* were done in Chimera (Pettersen et al., 2004). For item *b* sequences were aligned with the web tool MUSCLE (Edgar, 2004) and visualized with CPRISMA program 1.0 (Poveda-Cuevas, 2021) Regarding item *d*, the “ProteinVolume server” was employed (Chen and Makhatadze, 2015).

### 2.3. Coarse-grained model

The NS1 proteins are relatively large objects which imply several interacting sites in a computer simulation with available classical force fields. For instance, the NS1_DENV2_ homodimer has 704 amino acids (5604 atoms). The need to repeat the calculations for four different biological systems (NS1_ZIKV-UG_, NS1_ZIKV-BR_, NS1_DENV2_, and NS1_WNV_) at two oligomeric states (dimers and tetramers) and at several pH and salt regimes makes simplified molecular models quite appealing for this study. Moreover, the aim is also to investigate the driving forces behind self-association. This requires estimating the free energy of interactions [*βw*(*r*), where *β* is equal to 1/*k*_*B*_*T, k*_*B*_ *=* 1. 380 × 10^−23^ m^2^ kg s^-2^ K^-1^ is the Boltzmann constant, and *T* is the temperature in Kelvin] as a function of separation distances (*r*) between the centers of the proteins. It is well known that such measurements are very expensive in terms of computational time. Longer simulation runs are needed to ensure both that the most important parts of the configurational space have been properly sampled and that the histogram bins used to estimate *βw*(*r*) are well-populated within a reasonable number of observations to be statistically meaningful (Barroso da Silva et al., 2006; Engkvist and Karlström, 1996). These reasons together motivated us to invoke a constant-pH (CpH) coarse-grained (CG) model especially devised before to explore protein-protein interactions (Da Silva et al., 2016; Delboni and Da Silva, 2016; Giron et al., 2020; Mendonça et al., 2019). Such a simplified biophysical model with a reduced number of degrees of freedom and input parameters has already proved to be advantageous and suitable for this kind of purpose at a lower (not necessarily cheaper) and feasible computational cost (Da Silva and Dias, 2017). Accordingly, amino acids are converted into a set of charged Lennard-Jones (LJ) spheres (beads) with radii *R*_*ai*_ and valences *z*_*ai*_. The resulting mesoscopic structures are treated as rigid bodies (*i*.*e*., bond lengths, angles, and dihedral angles are kept fixed). The size and density of each of these charged LJ spheres were defined following the parameters given by Persson et al. (2010). The valences of the titratable residues are calculated as a function of pH during the simulation runs with the fast proton titration scheme (FPTS) (Da Silva and Dias, 2017; Da Silva and MacKernan, 2017; Teixeira et al., 2010). Intra and interchain charged amino acids can interfere in the acid-base properties of any other ionizable group as expected. Non-ionizable residues are not affected by the FPTS scheme. They are kept neutral at all times.

During the simulation runs the NS1 proteins are placed in an electroneutral cylinder with radius *r*_*cyl*_ (all systems = 350 Å) and length *l*_*cyl*_ (monomer-monomer = 200 Å, and dimer-dimer = 350 Å), where they can titrate, rotate in any direction, and translate back and forward along one axis. This strategy has been recently named FORTE (**F**ast c**O**arse-grained p**R**otein-pro**T**ein mod**E**l) (Neamtu et al., 2022). A detailed discussion of this theoretical approach, its pros and cons were given in previous works (Da Silva and Dias, 2017; Da Silva and MacKernan, 2017; Teixeira et al., 2010). Lipids and sugars are not present in the model. For the dimerization process, two single monomers were placed in the simulation box. In an analogous manner, two dimers were present in the simulation box to study the tetramerization process. Water molecules are effectively modeled by a static dielectric constant ϵ_*s*_ equals to 78.7 (at 298 K). The main effect of counterions and added salt particles is given by an electrostatic screening term [*exp*(− κ*r*_*ij*_)], where κ is the inverse modified Debye length [more details are given in the refs. (Da Silva and Dias, 2017; Da Silva and MacKernan, 2017)], and *r*_*ij*_ is the inter amino acid separation distance (Teixeira et al., 2010). A sketch representing the model is shown in Figure 2.

**Figure 2.**
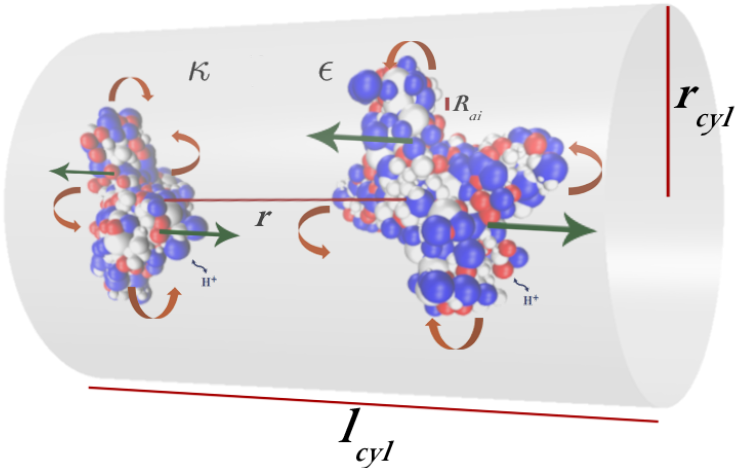
Schematic representation of the model cell used in this study. Two proteins build up by a collection of charged LJ spheres of radii (*R*_*ai*_) and valence *z*_*ai*_ mimicking amino acids. The simulation “box” is an electroneutral open cylindrical cell of radius *r*_*cyl*_ and height *l*_*cyl*_. The solvent is represented by its static dielectric constant ϵ_*s*_. Counter-ions and added salt particles are represented by Debye’s term κ. Acidic, basic, and neutral residues are represented in blue, red, and white respectively. See the text for more details.

The effective configurational Hamiltonian of the system for a given configuration [*U*({*r*_*k*_})] is calculated as,

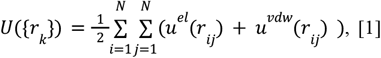

where {*r*_*k*_} are the positions of the amino acids, *N* is the total number of amino acids, [*u*^*el*^ (*r*_*ij*_)] is the Coulombic contribution, [*u*^*vdw*^(*r*_*ij*_)] is the LJ term, and *r*_*ij*_ is the separation distance between amino acids *i* and *j*.

The potential energy *u*^*el*^ (*r*_*ij*_) describes the electrostatic interaction between any two ionizable groups of valences *z*_*i*_ and *z*_*j*_:

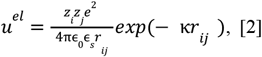

Where ϵ_0_ is the dielectric constant of the vacuum (*=* 8. 854 × 10^−12^C^2^/Nm^2^) and *e* is the elementary charge (*=* 1. 602 × 10^−19^ C).

Using different radii for the amino-acids, the van der Waals (vdw) interactions partially incorporating the non-specific contributions of the hydrophobic effect and the Pauli exclusion principle are effectively calculated with the *u*^*vdw*^ (*r*_*ij*_) potential energy as given by,

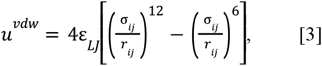

where ε_*LJ*_ is an adjustable parameter that defines the depth of the potential well. It can be tuned up to favor the complexation or down to result in a lower attraction depending on the simulated system. Here, we assumed a more or less universal value for protein studies where ε_*LJ*_ is equal to 0.124 kJ/mol as suggested by Persson and co-authors (Persson et al., 2010). This should correspond to a Hamaker constant of ca. 9 *k*_*B*_*T* for an amino acid pair (Lund and Jönsson, 2003). The variable σ_*ij*_[= *R*_*ai*_ + *R*_*ai*_)/2] is related to the size of beads (*R*_*ai*_ and *R*_*ai*_). These values for each amino acid were taken from the study by Da Silva et al. (2016). Larger amino acids should strongly attract others which results in a reasonable description of macromolecular hydrophobic moments (Eisenberg et al., 1982; Lund and Jönsson, 2003).

### 2.4. Constant-pH Monte Carlo simulations

The molecular systems were simulated at different pH solutions and salt concentrations by Monte Carlo (MC) simulations using the Metropolis algorithm (Metropolis et al., 1953). Free energy derivatives were computed as a function of the separation distance for both monomer-monomer and dimer-dimer complexations. For the sake of convenience, several sets of simulations in the NVT ensemble were performed for all viruses/strains:

1. **Set A:** A single NS1 protein (monomeric state) in NaCl solution (150 mM). This set was used to provide the main physical-chemical properties of the proteins;
2. **Set B:** Same as set A for the dimeric state;
3. **Set C:** Two NS1 monomers in salt solution (150 mM) during the dimerization processes (NS1_flavivirus_m_ + NS1_flavivirus_m_ → NS1_flavivirus_mm_). Here, both macromolecules can translate, rotate, and titrate to sample the complexation process. These simulations were employed to provide *βw*(*r*) for the dimerization. Note that the resulting dimer was labeled with the subscript “mm”;
4. **Set D:** Same as set C for the tetramerization processes (NS1_flavivirus_d_ + NS1_flavivirus_d_ → NS1_flavivirus_dd_). NS1_flavivirus_d_ is the dimer as given by the experimental structure. Note that here the resulting tetrameter was labeled with the subscript “dd”;
5. **Set E:** Same as set D for a hypothetical system with all amino acids assumed to be constantly neutral during the simulation run. This artificial system was solely used to access the free energy contributions exclusively from the van der Waals terms excluding the electrostatic interactions;
6. **Set F:** Same as set D for lower ionic strength (5 mM).

For sets A and B, the equilibration and production phases were performed using 10^6^ and 10^8^ MC cycles, respectively. Only titration was included as an MC movement in these runs. The FPTS was employed at a different pH condition from 0.1 to 14 every 0.1 pH unit to compute the corresponding protein net charge number (*Z*_*P*_), the charge regulation capacity (*C*_*P*_), and the dipole moment number (*μ*_*P*_) as previously proposed (Delboni and Da Silva, 2016; Jonsson and Lund, 2007; Lunkad et al., 2022; Poveda-Cuevas et al., 2018). For the other simulation sets where the focus was on the complexation process, much longer and quite expensive (in terms of CPU time) simulations were necessary. At least 1.0 × 10^10^ and 2.0 × 10^10^ MC steps were carried out for the dimerization and tetramerization processes, respectively, after the equilibration phase. Due to the high computational cost of these simulations, a bit shorter runs were executed for sets E and F. In these runs (*i*.*e*. E and F), 2.6 × 10^8^ MC and 9.0 × 10^8^ steps were employed, respectively. In all sets, three replicates were carried out for each simulated system. They were also used to estimate mean values and standard deviations. All MC simulations were done through a modified version of the biomolecular simulation package Faunus (Lund et al., 2008).

The central aspects of the protein-protein complexation process were covered by sets C and D. Three different pH solutions were explored: 3, 7, and 8.5. These values correspond to the experimental conditions where dissociation and association were reported for NS1_DENV2_ (Falconar and Young, 1990). Previously electrophoresis assays and X-ray data have shown that the dimerization occurs at a pH interval from 3.5 to 8.5 (Akey et al., 2014; Brown et al., 2016; Falconar and Young, 1990; Flamand et al., 1999; Song et al., 2016; Xu et al., 2016). At pH 3, monomers were observed instead of the dimers (Falconar and Young, 1990). Few experimental information is available for the tetramerization. There are only two reported cases at pH 8 and 8.5 for NS1_DENV2_ and NS1_DENV1_, respectively (Flamand et al., 1999; Muller et al., 2012). Therefore, the choice of the pH regimes in this work was twofold: *a*) to validate the model capacity to reproduce experimental data (for NS1_DENV2_), and *b*) to apply it to other conditions and viral systems where there is a lack of experimental data (*e*.*g*. pH 7 and NS1_ZIKV-UG_dd_). To uncover the impact of the specific protein sequences, it is important to compare the behavior of NS1 proteins from NS1_ZIKV-UG_, NS1_ZIKV-BR_, NS1_DENV2_, and NS1_WNV_ on exactly the same experimental conditions. Other pH conditions were not simulated due to the already high CPU costs of these three selected conditions. For each single MC step, the computing time was 0.06 seconds (processor Intel(R) Xeon(R) CPU E7-2870 @ 2.40GHz) for a dimer-dimer complexation study. This number grows too fast when multiplied by the needed ca 3 × 10^10^ steps and the replicates together additional tests per case and condition.

The last two sets (E and F) were complementary runs to provide the energy basis for the self-association of these NS1 proteins. Simulations with all ionizable groups artificially forced to keep zero their charges are used in set F to quantify the vdw contributions on each flavivirus NS1 system (in the absence of electrostatic interactions). Finally, set E was carried out at pH 7 to assess the role of a low salt condition (5 mM) on the tetramerization process.

### 2.5. Free energy of interactions

The main result from sets C to F is *βw*(*r*). During each simulation run, it was recorded in histograms the frequency of each separation distance between the center of the proteins observed. The sampling was performed with a bin size equal to 1Å. These histograms were converted to radial distribution functions [*g*(*r*)]. Through a well-established mechanical-statistical relation, *g*(*r*) is related to the two-particle potential of mean force [*βw*(*r*) = −ln *g*(*r*)] that was used to calculate the corresponding free energy of interactions (McQuarrie, 1976).

### 2.6. Second virial coefficient

When a pair of proteins are immersed in an electrolyte solution, several physical interactions are acting on the system (*e*.*g*. repulsion due to the Pauli-exclusion principle, vdw, hydrophobic “interactions” and multipole-multipole electrostatic interactions). Practical estimation of their importance can be done involving the calculation of second virial coefficients (*B*_2_). Such quantity can also be obtained by means of light scattering (Velev et al., 1998), chromatography (Tessier et al., 2002), or membrane osmometry experiments (Pjura et al., 2000). Theoretically, *B*_2_ (often given in mol ml/g^2^) can be estimated following the canonical partition [*B*_2_ = *B*_*q*+*vdw*_ + *B*_*hs*_], where *B*_*q*+*vdw*_ represents the electrostatics and vdw contributions, and *B*_*hs*_ represents the excluded-volume (hard-sphere) term. Mathematically,

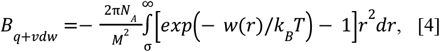

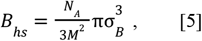

where *M* is the molecular weight (in g/mol) of proteins, *σ*_*B*_ is the sum of radii of the two proteins (in Å) and *N*_*A*_ is the Avogadro number (6.022×10^23^ mol^-1^). Equations 4 and 5 are numerically solved through the Simpson rule from *βw*(*r*) given by the MC simulation data and simple analytical math, respectively. Values for *B*_*hs*_ for the NS1 protein systems in the monomeric and dimeric state for each NS1_flavivirus_ are compiled in Table S1. This data does not depend on the simulation results as seen in Eq. 5. A positive *B*_2_ (total term) indicates an attractive behavior, whereas a negative value means a repulsion between the proteins (George and Wilson, 1994). The dependence on *r*^*2*^ in Equation 4 makes *B*_*q*+*vdw*_ largely affected by the statistical noises of *w(r)*. Much longer runs would be needed for a precise quantitative prediction of *B*_2_. Instead, due to the prohibitive CPU costs we focuses on the qualitative trends given by the calculated *B*_2_. The molecular weights (*Mw*) of the studied proteins were estimated in the web server tool “Compute pI/Mw” (Wilkins et al., 1999). The *Mw*’s in the monomeric states are 40078.36, 40097.47, 39939.47, and 39749.8 g/mol, for respectively, NS1_ZIKV-UG_, NS1_ZIKV-BR_, NS1_DENV2_, and NS1_WNV_. For all NS1_flavivirus_ in the dimeric state these, *Mw*’s were doubled.

## 3. RESULTS AND DISCUSSION

The systems studied in the present paper cover different ranges of viral time-scale evolution, a short-range for the two ZIKV strains (Gong et al., 2016), and a long-range between ZIKV, DENV2, and WNV (Moureau et al., 2015; Pettersson and Fiz-Palacios, 2014). This is illustrated by the identity percentages between the four NS1_flavivirus_ as reported in Table S2. The NS1_ZIKV_ strains present 97% identity, while the remaining pairs have an average identity percentage equivalent to 55.2% when compared. The flavivirus NS1 protein is a multifaceted molecule often referred to as a promiscuous biological system due to its ability to interact with different partners (Muller and Young, 2013; Watterson et al., 2016). This can be advantageous from the point of view of viruses since the cellular machinery is integrated by several macromolecules and biological media that should be altered to produce the host-cell programming. Electrostatic forces have an impact on these interactions as it has already been seen in both experiments and computer simulations (Brown et al., 2016; Poveda-Cuevas et al., 2020, 2018; Song et al., 2016; Xu et al., 2016). A considerably high number of titratable sites (see Figure 1 and S1) is observed for both monomers and dimers. Table S3 lists the number of ionizable residues for each flavivirus NS1 protein at the monomeric state. Note that these four viral systems have a different number of titratable groups. The two NS1_ZIKV_ have a similar number of basic (66 and 67, for NS1_ZIKV-UG_ and NS1_ZIKV-BR_, respectively) and acidic residues (51 and 50, for NS1_ZIKV-UG_ and NS1_ZIKV-BR_, respectively) because they are phylogenetically closer while NS1_DENV2_ and NS1_WNV_ exhibit a larger difference because they belong to other flavivirus species. In terms of their three-dimensional structures, RMSD calculations among the C_α_ for all possible pairs of NS1 dimers show some structural differences among them. The most and least similar NS1 dimers correspond to the pairs NS1_ZIKV-UG_-NS1_DENV2_ (0.492 Å) and NS1_ZIKV-BR_-NS1_WNV_ (0.765 Å), respectively (see Table S4). A detailed view when comparing the C_α_s of the NS1 structures per residue shows that regions with the greatest structural variations as measured by the RMSD comprise the loop zones between residues 115-122 (see Figure S1). These two features (*i*.*e*. number/composition of titratable residues and a specific three-dimensional structure for each NS1 protein with a different spatial arrangement of the ionizable residues) should have an impact on electrostatic properties and the biomolecular interactions of these macromolecules. Consequently, physical-chemical variables such as pH are important in the molecular mechanisms that control the formation of NS1 complexes and impact the virus life cycle.

Free energies of interactions [*βw*(*r*)] and second virial coefficients (*B*_*2*_) were employed to desiccate the self-association features of the NS1_ZIKV-UG_, NS1_ZIKV-BR_, NS1_DENV2_, and NS1_WNV_ at different pH conditions: 3 (acidic regime), 7 (neutral/physiological regime), and 8.5 (basic regime). These free energies were computed assuming an ionic strength of 150 mM of NaCl. On the basis of these physicochemical conditions, this section focuses four aspects: *i*) the validation of our theoretical model using the available experimental information, *ii*) a report of the specificities of these viruses for both dimerization (NS1_flavivirus_m_ + NS1_flavivirus_m_ → NS1_flavivirus_mm_) and tetramerization (NS1_flavivirus_d_ + NS1_flavivirus_d_ → NS1_flavivirus_dd_) processes, *iii*) the examination of the discrepancies at the inter-(using the four NS1 flavivirus proteins) or intraspecific level (focusing in the two NS1_ZIKV_ proteins), and *iv*) the analysis of the physical forces that govern the self-association and their possible biological implications. The *βw*(*r*) for both NS1_flavivirus_mm_ and NS1_flavivirus_dd_ for all four flaviviruses were calculated through CpH MC. Figure 3 displays the outcomes of these simulations for all studied pH conditions and the two oligomeric states.

**Figure 3.**
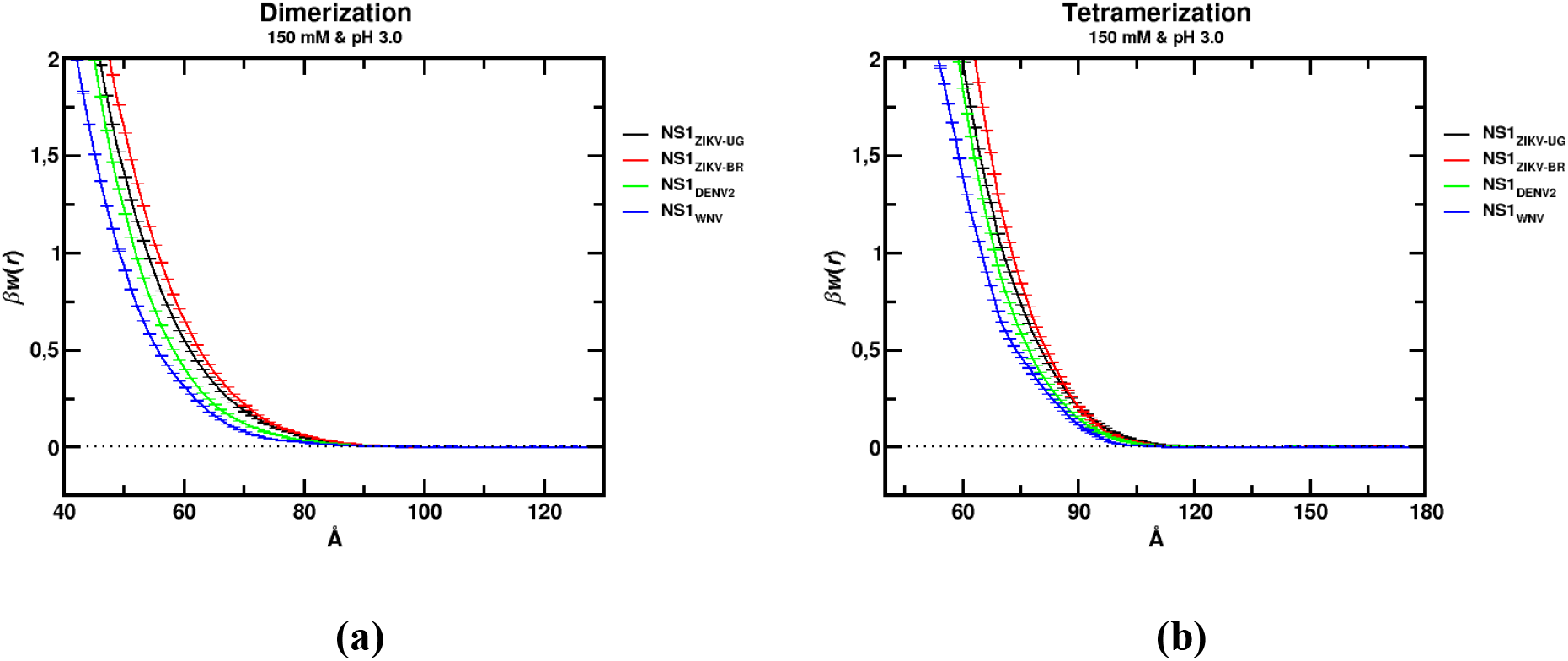
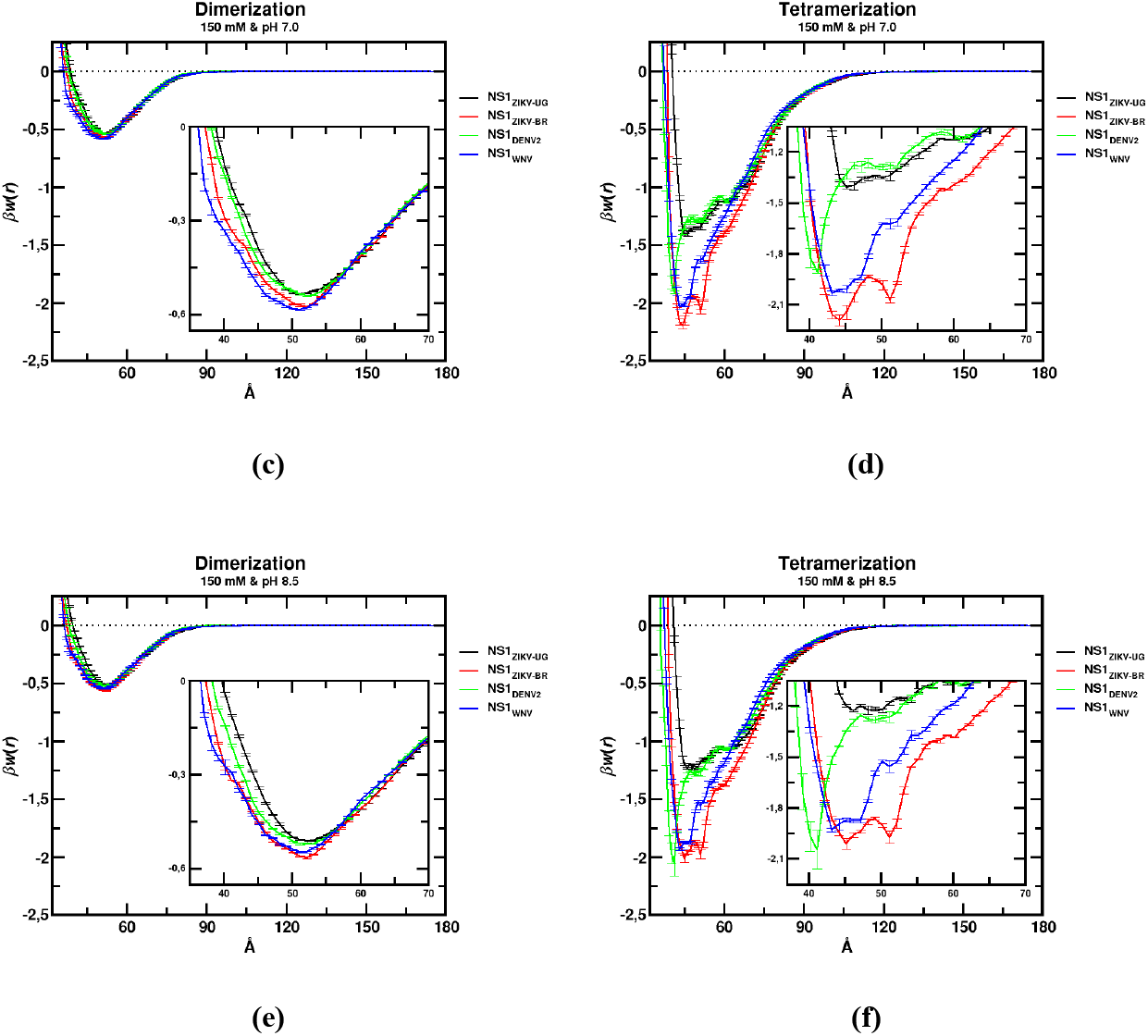
Free energy of interactions as a function of the center-center separation distance for NS1 homoassociation. Data for the dimerization and the tetramerization processes of four flaviviruses at different pH solutions and 150 mM of NaCl. The plots show the average data with the estimated standard deviations based on three replicate runs for each studied system. These data correspond to the results from simulation sets C (NS1_flavivirus_mm_) and D (NS1_flavivirus_dd_).

### 3.1. Validation of the computational model

A common procedure to validate computational simulations of biomolecules is to compare specific physical quantities predicted by the models with experimental data.(Cragnolini et al., 2015) The reproducibility of the experimental information by the theoretical model allows establishing its predictive capacity for subsequent novel applications (*e*.*g*., the prediction of the tetramerizations features that are not yet experimentally available in the literature). Although the model has been validated for other systems (Da Silva et al., 2016; Delboni and Da Silva, 2016; Giron et al., 2021, 2020; Neamtu et al., 2022), for this purpose and NS1 proteins, we have mainly used electrophoresis data by Falconar and Young (1990) for dimerizations of NS1_DENV2_ (*i*.*e*., NS1_DENV2_mm_). Data is available for the pH range of 2.2 to 7.4. It was found in this lab experiment that for situations above pH ∼3 the system forms stable dimers, whereas below pH ∼3 a monomeric state prevails. No electrophoresis data is available for the other three NS1 proteins and/or pH conditions. Some additional information can be extracted from structural data. For instance, combining the crystallography conditions where crystals for NS1_ZIKV-UG_mm_ and NS1_ZIKV-BR_mm_ (Brown et al., 2016; Xu et al., 2016) were obtained with electrophoresis data for NS1_DENV1_dd_ and NS1_DENV2_dd_ (Flamand et al., 1999; Muller et al., 2012) indicates that there are dimers or tetramers above pH 8. This was tested in our biophysical simulations. The *βw*(*r*) profiles for the dimerization process of NS1_DENV2_mm_ were calculated at pH values of 3, 7, and 8.5 (see, respectively, the green curves in Figures 3a, 3c, and 3e). Following a conventional thermodynamic criterion to identify repulsive or attractive systems [*i*.*e*., when *βw*(*r*) > 0 and *βw*(*r*) < 0, respectively] (McQuarrie, 1976), we observed that monomers of NS1_DENV2_mm_ form homodimers at pH 7 and 8.5, while they are unable to associate at pH 3 in agreement with the results of the electrophoretic experiment. For the NS1_ZIKV-UG_mm_ and NS1_ZIKV-BR_mm_ at pH 8.5 (see black and red curves in Figure 3e), our model predicts attractive profiles for both viral proteins. The same was found for NS1_DENV2_dd_ that experiences an attraction at pH 8.5 confirming it can self-associate at this condition (see Figure 3f). For all cases, our theoretical model is in good agreement with the available experimental data predicting that dimers and monomers can be found at the correct solution pH value. This allows us to further explore other new conditions in order to comprehend more about the NS1 self-association of these four viral systems (in particular the tetramerization process). It is important to mention that the model does not incorporate conformational changes that can occur under extreme pH conditions (*i*.*e*., unfolding at acidic or basic regimes far from the protein isoelectric point). Nevertheless, no discrepancy was observed for the predictions at the acidic pH in comparison with the experiments.

### 3.2. Differences at the oligomerization level

Next, we analyzed and compared the NS1 dimerization and tetramerization processes at the same physicochemical conditions for the four viral systems. We found that regardless of the type of virus, strain, or oligomerization state, the four systems are all repulsive at pH 3 as indicated by their estimated free energy of interactions (*βw*(*r*) > 0 - see Figures 3a and 3b). At this pH, the tetramerization is even more unfavorable compared to dimerization on the range of separation distances examined. Conversely, an attractive regime [*βw*(*r*) < 0] is observed for all NS1 proteins at pH 7 for both the dimerization (see Figure 3c) and tetramerization (see Figure 3d) processes. A higher well-depth is measured for the tetramerization in comparison with the dimer formation. For instance, using the minima values of the *βw*(*r*) potential well as a parameter for this analysis, the attraction is 4-fold larger for NS1_ZIKV-BR_dd_ than NS1_ZIKV-BR_mm_ [*βw*(*r*)_min_dd_ = -2.19±0.03 *k*_*B*_*T* at 44.2 Å and *βw*(*r*)_min_mm_ = -0.575±0.002 *k*_*B*_*T* at 52.2 Å, for NS1_ZIKV-BR_dd_ and NS1_ZIKV-BR_mm_, respectively]. This behavior suggests an important contribution from the van der Waals interactions for these complexes. The theoretical predictions for *βw*(*r*) at pH 7 and 8.5 are quite similar. These data can be seen in Figures 3e and 3f, respectively. The main difference is the slight tendency for the protein-protein association to be less attractive at pH 8.5 due to the increased Coulombic repulsion at this more basic condition [*e*.*g*., *Z*_*P*_s are -4.3 (pH 7) and -10.4 (pH 8.5) for NS1_ZIKV-UG_d_].

A complementary analysis can be done by means of second virial coefficients. They can be roughly estimated from the computed *βw*(*r*) as shown in Equation 4. The quantity *B*_*2*_ is interpreted similarly as for *βw*(*r*), that is, when a system undergoes an attractive regime, *B*_*2*_ < 0, while for a repulsive condition, *B*_*2*_ > 0. Figure 4 shows *B*_*2*_ as a function of pH for all studied systems (inter and intraspecific levels), oligomeric states, and physical-chemical conditions. The data for the formation of dimers and tetramers are given by the dashed and solid curves, respectively. It can also be easily seen in this plot how tetramerizations are more attractive than dimerizations for any pH solution. All solid lines (tetramerization) are always below the dashed ones (dimerization) for all systems. Even at pH 3, lower repulsive features are observed for the dimer-dimer interactions compared to the monomer-monomer association, in contrast to the data represented in Figure 3a and Figure 3b. For instance, *B*_*2*_ decreases from 1.90±0.02 ml mol × 10^−4^/g^2^ (for the dimer formation) to 1.022±0.006 ml mol × 10^−4^/g^2^ (for the tetramer formation) at pH 3 for NS1_WNV_. This change might be explained by a smaller repulsive excluded-volume contribution for the dimer-dimer association compared to monomer-monomer interaction (see Table S1).

**Figure 4.**
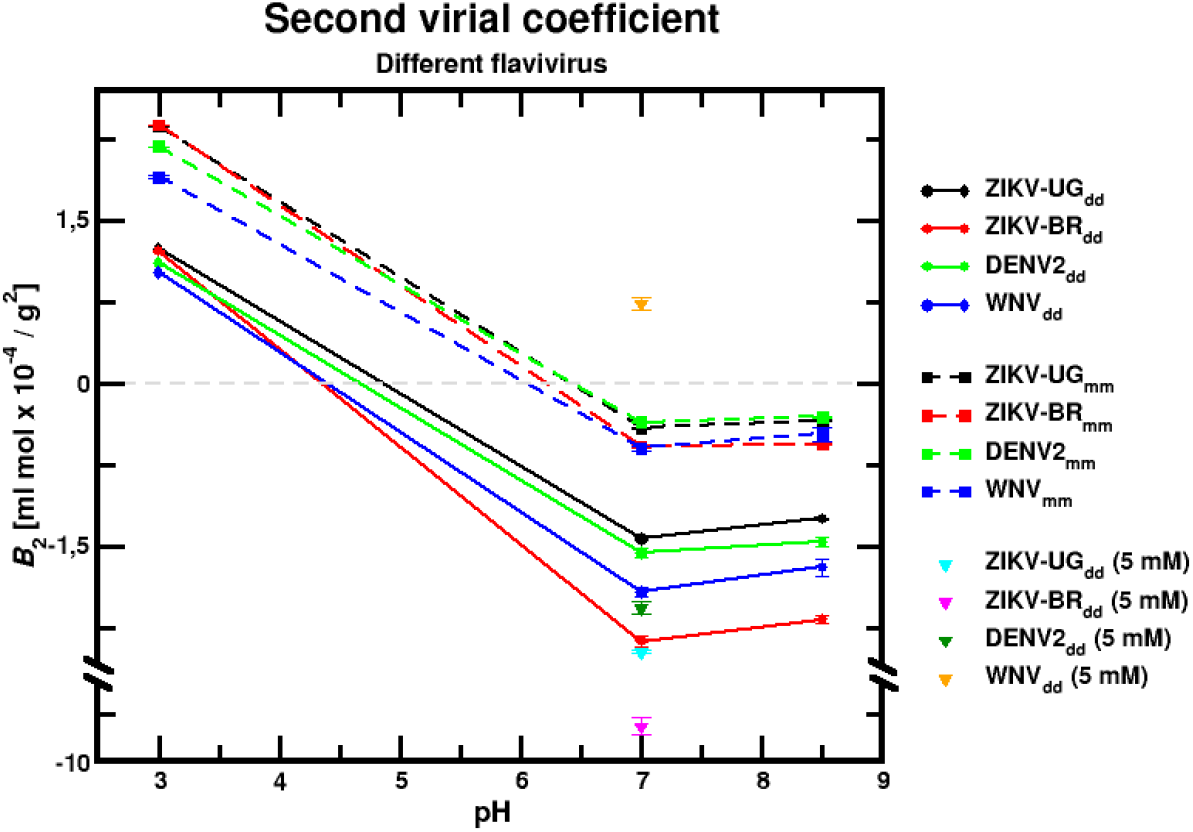
Second virial coefficient *B*_2_ as a function of pH for several flavivirus NS1 and two oligomeric states at different pH solutions. The plots show an average behavior estimated on three replicate runs. Dashed and continuous lines are used to guide the eyes for the formation of dimers and tetramers, respectively. All symbols connected by these lines represent data obtained at 150mM of NaCl. Symbols not connected by lines represent data obtained at a low salt condition (5mM of NaCl).

### 3.3. Differences at the inter and intraspecific levels

For the understanding of the specific molecular mechanisms that the NS1 protein might obey for each viral system, we compared their associative characteristics. In Figures 3a and 3b, we start with the comparison at pH 3 for their self-associations. For both oligomeric states, the four viral systems can be sorted out from the more repulsive behavior to the less one based on the minima values of their free energies of interactions: *βw*(*r*)_ZIKV-BR_ > *βw*(*r*)_ZIKV-UG_ > *βw*(*r*)_DENV2_ > *βw*(*r*)_WNV_. At physiological pH, the monomer-monomer association follows a different order: *βw*(*r*)_min_ZIKV-UG_mm_ > *βw*(*r*)_min_DENV2_mm_ > *βw*(*r*)_min_ZIKV-BR_mm_ > *βw*(*r*)_min_WNV_mm_ as seen in Figure 3c. The NS1 protein from the WNV is the system with the highest NS1-NS1 binding affinity. Yet, for the tetramer formation, there is an inversion between NS1_WNV_dd_ and NS1_ZIKV-BR_dd_: *βw*(*r*)_min_ZIKV-UG_dd_ > *βw*(*r*)_min_DENV2_dd_ > *βw*(*r*)_min_WNV_dd_ > *βw*(*r*)_min_ZIKV-BR_dd_ (see Figure 3d). The same trend is seen at pH 8.5 as illustrated in Figures 3e and 3f for the dimerization and the tetramerization processes, respectively. This result also implies that each system will have a different pH condition for the transition between a repulsive to an attractive regime. The connecting lines are drawn to guide the eyes in Figure 4. The minima values of the well-depth for these free energies of interactions are found at different separation distances. This is a direct consequence of the different sizes that each system has (see Table S5). Some NS1 proteins can come closer to their selfpartners than others. For instance, it is observed that the dimers of NS1_DENV_dd_ present a high frequency of interaction in a short distance range (notice in Figures 3d and 3f how the green curves are narrower at ∼40 Å).

The main features of this analysis can be summarized with the *B*_*2*_ data as given in Figure 4. For instance, for the formation of tetramers, from the less attractive regime to the more one, the systems are following this order: *B*_*2_*NS1_ZIKV-UG_dd_ > *B*_*2_*NS1_DENV2_dd_ > *B*_*2_*NS1_WNV_dd_ > *B*_*2_*NS1_ZIKV-BR_dd_. This can be seen at both pH 7 and 8.5 where the systems are under an attractive regime condition. In all the simulated conditions, lipids and sugars were absent from the system indicating that they are not necessary for these self-associations.

It is interesting to see how more phylogenetically related proteins such as NS1_ZIKV_UG_ and NS1_ZIKV-BR_ (97% of sequence identity among them) behave in a discrepant way. For example, at pH 7 (see Figure 3d), the smaller value for *βw*(*r*) was found for NS1_ZIKV-BR_dd_ (∼2 *k*_*B*_*T* at *r*∼44.5 Å) when compared to NS1_ZIKV-UG_dd_ (∼1.4 *k*_*B*_*T* at *r*∼45.5 Å). This means a difference of 0.6*k*_*B*_*T*. Conversely, the same comparison for NS1_ZIKV-BR_dd_ and NS1_WNV_dd_ gives a smaller difference of 0.2*k*_*B*_*T* despite their lower sequence identity (56%). This shows an interplay of the contributions determined at the linear sequence and three-dimensional structural levels. It also reveals an important diversity in the molecular mechanisms of these viral proteins with direct consequences on their virulences.

### 3.4. Physicochemical features

The evaluation of the main physicochemical aspects that govern these systems can shed light to understand the physical reasons for the differences discussed above (Da Silva et al., 2016; Delboni and Da Silva, 2016; Giron et al., 2021, 2020; Poveda-Cuevas et al., 2018). Therefore, we next characterized the main physical-chemical quantities (*Z*_*P*_, *C*_*P*_, and *μ*_*P*_) of NS1_ZIKV-UG_, NS1_ZIKV-BR_, NS1_DENV2_, and NS1_WNV_ for both monomeric and dimeric states at physiological salt concentration and pH values from 0 to 14. Based on a previous study (Poveda-Cuevas et al., 2018), the total protein charge number was identified as the central outcome for the present work due to the electrostatic screening of the high order terms. This physical quantity seems especially important in viral proteins (Giron et al., 2021; Poveda-Cuevas et al., 2020, 2018; Yoshida et al., 2019). Table 1 provided the main data for pH conditions of 3, 7, and 8.5. For the sake of completeness, full titration plots for the two oligomeric states and the four NS1 flaviviruses are also shown in Figure S2. Monomers can be highly charged at very acidic and basic regimes with *Z*_*P*_ ranging from ca. 53 to -53 (see Figure S2a). These protein net charges are relatively doubled in the dimeric state due to the presence of the second chain (see Figure S2b) that contributes to the charge regulation. Regarding the *C*_*P*_s of monomers (see Figure S2c) and dimers (see Figure S2d), we observed two peaks at roughly pH 3.7 and 11.6. Note that the *C*_*P*_ values are 2-fold large for NS1s in the dimeric state when compared to monomers (*e*.*g*., the *C*_*P*_s are 7.5 and 14.4 at pH 4.0 for, respectively, NS1_DENV_m_ and NS1_DENV_d_). This means that the second monomer in the dimeric NS1 is contributing to increasing *C*_*P*_. As explained before by Jonsson and Lund (2007) the reason for these two peaks has a direct relationship with the number of titratable sites and their p*K*_*a*_s values. Data for *μ*_*P*_ is given for the monomers and dimers, respectively, in Figures S2e and S2f. At pHs conditions closer to 7, NS1_ZIKV-UG_ and NS1_ZIKV-BR_ in the dimeric states have a particular characteristic presenting higher dipole values than the remaining NS1_flavivirus_. These observations indicate that charge regulation and dipole interactions (Barroso da Silva et al., 2006; Da Silva and Jönsson, 2009; Lunkad et al., 2022) can play a more important and specific role for each viral system when the salt concentration is sufficiently low.

**Table 1.** Protein charge number (*Z*_*P*_) for NS1_ZIKV-UG_ (PDB id 5K6K), NS1_ZIKV-BR_ (PDB id 5GS6), NS1_DENV2_ (PDB id 4O6B), and NS1_WNV_ (PDB id 4O6D). Data correspond to monomers (m) and dimers (d) at pH 3 (acid regime), 7 (neutral regime), and 8.5 (basic regime). The pIs for both oligomeric states are 6.7_pI_m_/6.6_pI_d_, 7.1_pI_m_/7.0_pI_d_, 6.8_pI_m_/6.8_pI_d_, and 5.8_pI_m_/5.8_pì_d_, for respectively NS1_ZIKV-UG_, NS1_ZIKV-BR_, NS1_DENV2_, and NS1_WNV_. Data correspond to simulation sets A and B.

| pH | ZIKV-UG |  | ZIKV-BR |  | DENV2 |  | WNV |  |
| --- | --- | --- | --- | --- | --- | --- | --- | --- |
| | $Z_{p\_m}$ | $Z_{p\_d}$ | $Z_{p\_m}$ | $Z_{p\_d}$ | $Z_{p\_m}$ | $Z_{p\_d}$ | $Z_{p\_m}$ | $Z_{p\_d}$ |
| 3 | 43.7 | 83.5 | 45.1 | 86.2 | 40.0 | 77.6 | 38.2 | 73.7 |
| 7 | -1.8 | -4.3 | 0.2 | -0.5 | -0.9 | -2.0 | -5.2 | -10.6 |
| 8.5 | -5.1 | -10.4 | -3.2 | -6.9 | -3.6 | -7.2 | -7.7 | -15.5 |

We noticed a correlation between the total protein net charges and the minima values of *βw*(*r*) for both NS1_flavivirus_mm_ and NS1_flavivirus_dd_ at pH 3. That is, as the protein net charge is increased the protein-protein free energy of interaction becomes more repulsive. For instance, compare the blue (NS1_WNV_), green (NS1_DENV2_), black (NS1_ZIKV-UG_), and red (NS1_ZIKV-BR_) curves given in Figures 3a and 3b with the protein net charges given in Table 1. The protein net charges follow the same order (*Z*_*P*_WNV_ < *Z*_*P*_DENV2_ < *Z*_*P*_ZIKV-UG_ < *Z*_*P*_ZIKV-BR_) indicating that charge-charge interactions dominate this situation in both the monomer and dimers states. At physiological or basic pHs, this behavior is not clearly observed, namely, we do not see that NS1_WNV_dd_ as the most repulsive system among the remaining NS1 despite having the higher protein net charge. This may be an indication that van der Waals forces are dominating the self-association at these pH solutions. Electrostatic interactions are probably screened out by physiological salt conditions which allow the vdw interactions to overcome them at these pH conditions. Non-Coulombic contributions such as those arising from the multipole-multipole interactions and the charge regulation mechanism are even more severely affected by salt (Barroso da Silva et al., 2006). Thus, in certain conditions, it has been experimentally shown that the oligomerization processes of some viral proteins with the same net charge can be driven by vdw interactions or the hydrophobic effect (Kegel and van der Schoot, 2004; Thuman-Commike et al., 1998).

The contributions of the vdw interactions were further explored by means of an artificial man-created model where electrostatic interactions were removed from the system. That is, all the charges of the titratable groups were artificially kept equal to zero during dimer-dimer complexation runs at physiological salt concentration. The corresponding *βw*(*r*) plots for this situation are given in Figure S3 (red lines). For the sake of comparison, *βw*(*r*) for the same system at pH 7 are included in this figure as solid black lines. The results suggest that vdw forces dominate the complexation process at this pH condition in contrast to what was observed before at the acidic pH condition. The protein net charges are not sufficiently high to produce a repulsion as it is observed at pH 3. Moreover, since the vdw contributions have a direct relationship with the dimension of the systems (as could be seen in Equation 3), they found differences between the two oligomeric states are largely explained by the protein sizes (the volumes of the monomers and dimers are equivalent to ∼4.8 × 10^4^Å^3^ and ∼9.7 × 10^4^Å^3^, respectively). Some specific amino acids can also prevent the proteins from coming closer. For example, residues 101-135 are localized in the wing domain and have high thermal fluctuation (see Figure S1). They could easily contribute to a steric hindrance during the protein-protein assembly. Previous computational studies have also shown that these flexible regions can play a particular role for a specific NS1_flavivirus_ (Gonçalves et al., 2019; Menezes et al., 2020; Poveda-Cuevas et al., 2021).

The understanding of the balance between electrostatics and vdw interactions at different conditions can be complemented by investigating a low salt condition where the electrostatic screening is reduced. Calculations were performed at 5mM of NaCl and pH 7 for the dimer-dimer homoassociation (see Figure S4). Most systems (NS1_ZIKV-UG_dd_, NS1_ZIKV-BR_dd_, and NS1_DENV2_dd_) are more attractive at the low salt concentration regime: a) NS1_ZIKV-UG_dd_ (ca. -1.8_5mM_ < -1.4_150mM_ *k*_*B*_*T*), b) NS1_ZIKV-BR_dd_ (ca -4.2_5mM_ < -2_150mM_ *k*_*B*_*T*), and c) NS1_DENV2_dd_ (ca -2.2_5mM_ < − 1.9_150mM_ *k*_*B*_*T*). Conversely, NS1_WNV_dd_ shows a less attractive characteristic (ca -1.5_5mM_ > -2.0_150mM_ *k*_*B*_*T*). These trends were already included in Figure 4 showing the same behavior.

### 3.5. Biological relevance

NS1 is known both to interact with the immune system (Biering et al., 2021; Edeling et al., 2014; Hertz et al., 2017) and to increase the permeability of human endothelial cells from the lung, dermis, umbilical vein, brain, and liver (Puerta-Guardo et al., 2019). As shown through *in vitro* and *in vivo* experiments by Puerta-Guardo et al., (2019) the NS1 from a particular flavivirus is tissue-specific which can consequently produce particular physiopathology. Assuming physiological conditions of pH and salt (*i*.*e*., regimes where several biological processes happen), our computational outcomes show that at the molecular scale, the free energy of the NS1 oligomerization from four flaviviruses present meaningful discrepancies.

Given that each flavivirus has different stabilities (especially when tetramers are formed), we can hypothesize that the differential virulence has a direct contribution from the amino acid composition of each NS1 that, at the same time, impacts their structures, the vdw (to a great extent particularly at the physiological condition) and electrostatic interactions during the complexation process. Surprisingly, these discrepancies are not limited to the *inter*specific level, but also to the intraspecific level, showing that the versatility of NS1 can be extended to flavivirus variants. Furthermore, it is reasonable to assume that the oligomeric stability is crucial for subsequent interactions with other biomolecules that must occur. Thus, each flavivirus having a differential free energy profile could have a relation with the particular tropism expressed by each one or the specific organ affinity found by laboratory assays (Biering et al., 2021; Puerta-Guardo et al., 2019).

As mentioned earlier, regions 101-135 seem to be important for the biological function of NS1_flavivirus_. Previous studies have shown that this region is immunodominant and may work to inhibit the NS1 activity and pathogenicity via antibodies (Lai et al., 2017). Although these protein regions may be related to interactions with other macromolecules such as GAGs or membranes (Ci et al., 2021, 2020; Puerta-Guardo et al., 2019), we suppose that they may also be directly involved in the protein-protein NS1 assembly. This implies that these regions can work as a starting point to selectively inhibit oligomer association of different flavivirus species or strains. Finally, the detection of hotspots [*i*.*e*. regions or residues that can contribute significantly to protein-protein interaction as observed in other studies (Gonçalves et al., 2019; Persson et al., 2010)] for NS1 oligomerization gives important clues for the vaccine development, drugs, or therapies against flavivirus infections.

## 4. CONCLUSIONS

The dimerization and tetramerization processes for NS1_ZIKV-UG_, NS1_ZIKV-BR_, NS1_DENV2_, and NS1_WNV_ were examined through free energy derivatives at three different pH solutions (that is, 3, 7, and 8.5) and physiological salt concentration. As an essential step, the theoretical modeling was validated by comparing the available experimental data and free energy measurements for dimerizations and tetramerizations processes. Our outcomes confirmed the reliability of the used CG model also for this biological system. Complexation data for other flaviviruses and different experimental conditions that were lacking in the literature were provided. Our results show that these NS1 proteins can form both dimers and tetramers under conditions near physiological pH. The dimer-dimer interactions were found more stable than the monomer-monomer association as expected.

pH plays an important role mainly controlling the regimes where van der Waals interactions govern the NS1 self-associations. Two regimes could be detected that are common for all studied NS1 systems: *a)* a repulsive one, at pH 3, dominated by repulsive charge-charge interactions, and an attractive condition, at pHs 7 and 8.5, due to the vdw interactions to be able to overcome the electrostatic contributions. The number of additional peaks around the minima values of the free energies also suggests a complex energy landscape with steric contributions by the structural features of each system.

In previous studies, a probable mechanism that could explain the tetramerization and eventual hexamerization of NS1 has been hypothesized, where the interaction NS1-membrane is an essential step to produce stable assemblies among NS1 dimers during the secretion process (Gutsche et al., 2011). Nevertheless, despite the reduced number of parameters of our computational model, we have shown that the dimer-dimer interaction of NS1 can also occur in the *absence* of lipid and sugar molecules. Stable tetramers were formed in a pure electrolyte medium. Moreover, it was hypothesized that tetramers have a higher probability to interact with the immune system when compared to dimers or monomers because their greater stability facilitates the recognition of immunoglobulins.

Finally, this set of information gives new clues about the molecular mechanisms that drive the self-association of NS1 proteins. This could be beneficial for the development of drugs based on oligomerization inhibition. This study also provides a complementary view on the discussion related to flavivirus virulence and differential cellular tropism expressed at the inter and intraspecific levels. Despite the similarity at the protein sequence level, we found that each flavivirus has a well-characteristic NS1 protein-protein interaction profile.

## 5. ACKNOWLEDGMENTS

This work has been supported in part by the “Fundação de Amparo à Pesquisa do Estado de São Paulo” [Fapesp 2015/16116-3 and 2020/07158-2 (F.L.B.d.S.)], the Conselho Nacional de Desenvolvimento Científico e Tecnológico (CNPq) [CNPq 305393/2020-0 (F.L.B.d.S.)], the Swedish National Infrastructure for Computing [SNIC 2020/14-38 and SNIC 2021/3-6 (F.L.B.d.S.)] and the “Coordenação de Aperfeiçoamento de Pessoal de Nível Superior”—CAPES (S.A.P.C.). F.L.B.d.S. and S.A.P.C. acknowledge the support of the University of São Paulo (USP) through the NAP-CatSinQ (Research Core in Catalysis and Chemical Synthesis) and the computing hours at HPC/USP. The authors also would like to thank CINES (Centre Informatique National de l’Enseignement Supérieur) for computational resources (project: A0060710878). F.L.B.d.S. and C.E. acknowledge the mobility support from the joint Université Sorbonne Paris Cité-USP Research program. The authors also gratefully acknowledge the computing time granted by the John von Neumann Institute for Computing (NIC) and provided on the supercomputer JURECA at Jülich Supercomputing Centre (JSC) - project C210400511.

## Author Contributions

S.A.P.C., F.L.B.d.S., and C.E., performed simulations, analyzed data, and wrote the paper. F.L.B.d.S. and C.E. designed and supervised the research.

