## Supplementary material for "Self-association features of NS1 proteins from different flaviviruses": SI

[a]Universidade de São Paulo, Programa Interunidades em Bioinformática, Rua do Matão, 1010, BR-05508-090 São Paulo, São Paulo, Brazil.

[b]Universidade de São Paulo, Departamento de Ciências Biomoleculares, Faculdade de Ciências Farmacêuticas de Ribeirão Preto, Av. do Café, s/no-Campus da USP, BR-14040-903 Ribeirão Preto, São Paulo, Brazil.

[c] Department of Chemical and Biomolecular Engineering, North Carolina State University, Raleigh, NC 27695, United States.

[d]Université Paris Cité, Biologie Intégrée du Globule Rouge, Equipe 2, INSERM, F-75015 Paris, France.

[e]University of São Paulo and Université de Paris International Laboratory in Structural Bioinformatics, Faculdade de Ciências Farmacêuticas de Ribeirão Preto, Av. do Café, s/no-Campus da USP, Bloco B, BR-14040-903 Ribeirão Preto, São Paulo, Brazil.

* Corresponding Author


### SUPPLEMENTARY MATERIAL

**Table S1.** The excluded-volume (hard-sphere) term for the second virial coefficient. $B_{hs}$ (given in ml mol x $10^{-4}/g^2$) is calculated for the NS1 proteins at the monomeric and dimeric states using Equation 5. Parameters are given in the text and in Table S5.

| specie/strain | Monomers | Dimers |
|---|---|---|
| ZIKV-UG | 0.6182 | 0.3425 |
| ZIKV-BR | 0.5515 | 0.2871 |
| DENV2 | 0.7874 | 0.2893 |
| WNV | 0.6425 | 0.2921 |

**Table S2.** Percentage of sequence identities between pairs of NS1 flaviviruses proteins. Identities were calculated for all possible comparisons among the four NS1 flavivirus [PDB ids 5K6K ($NS1_{ZIKV\text{-}UG}$), 5GS6 ($NS1_{ZIKV\text{-}BR}$), 4O6B ($NS1_{DENV2}$), and 4O6D ($NS1_{WNV}$)]. Data is given in percentage. Values are calculated using Chimera (Pettersen et al., 2004).

| Identity for all possible NS1 pairs | | | | |
|---|---|---|---|---|
| specie/strain | ZIKV-UG | ZIKV-BR | DENV2 | WNV |
| ZIKV-UG | 100 | | | |
| ZIKV-BR | 97.4 | 100 | | |
| DENV2 | 54.8 | 53.8 | 100 | |
| WNV | 56.3 | 56.0 | 55.1 | 100 |

**Table S3.** The number of ionizable residues for four flavivirus NS1 proteins at the monomeric state. The numbers of each residue were calculated from the PDB ids 5K6K ($NS1_{ZIKV\text{-}UG}$), 5GS6 ($NS1_{ZIKV\text{-}BR}$), 4O6B ($NS1_{DENV2}$), and 4O6D ($NS1_{WNV}$).

| NS1 | pI | Asp | Glu | Tyr | His | Lys | Arg | Total acid | Total basic | Total |
|---|---|---|---|---|---|---|---|---|---|---|
| ZIKV-UG | 6.7 | 15 | 36 | 9 | 11 | 21 | 25 | 51 | 66 | 117 |
| ZIKV-BR | 7.1 | 16 | 34 | 8 | 12 | 22 | 25 | 50 | 67 | 117 |
| DENV2 | 6.8 | 15 | 29 | 8 | 10 | 28 | 13 | 44 | 59 | 103 |
| WNV | 5.8 | 19 | 30 | 9 | 8 | 21 | 21 | 49 | 59 | 108 |

**Table S4.** Structural differences among pairs of NS1 flaviviruses proteins. RMSDs were calculated between $C_{\alpha}$s of the NS1 dimers for all possible comparisons among the four NS1 flaviviruses [PDB ids 5K6K ($NS1_{ZIKV\text{-}UG}$), 5GS6 ($NS1_{ZIKV\text{-}BR}$), 4O6B ($NS1_{DENV2}$), and 4O6D ($NS1_{WNV}$)]. Data are given in Å. See the main text for more details.

| RMSD for all possible NS1 pairs | | | | |
|---|---|---|---|---|
| specie/strain | ZIKV-UG | ZIKV-BR | DENV2 | WNV |
| ZIKV-UG | 0 | | | |
| ZIKV-BR | 0.742 | 0 | | |
| DENV2 | 0.492 | 0.708 | 0 | |
| WNV | 0.613 | 0.765 | 0.598 | 0 |

**Table S5.** Molecular volumes for the NS1 proteins at the monomeric and dimeric states. Data is given in Å$^3$. The values were calculated at "ProteinVolume server" (Chen and Makhatadze, 2015) for $NS1_{ZIKV\text{-}UG}$ (PDB id 5K6K), $NS1_{ZIKV\text{-}BR}$ (PDB id 5GS6), $NS1_{DENV2}$ (PDB id 4O6B), and $NS1_{WNV}$ (PDB id 4O6D).

| specie/strain | Monomers | Dimers |
|---|---|---|
| ZIKV-UG | 47811.0 | 97222.3 |
| ZIKV-BR | 48182.5 | 97751.6 |
| DENV2 | 47332.1 | 96414.6 |
| WNV | 47687.9 | 96773.0 |

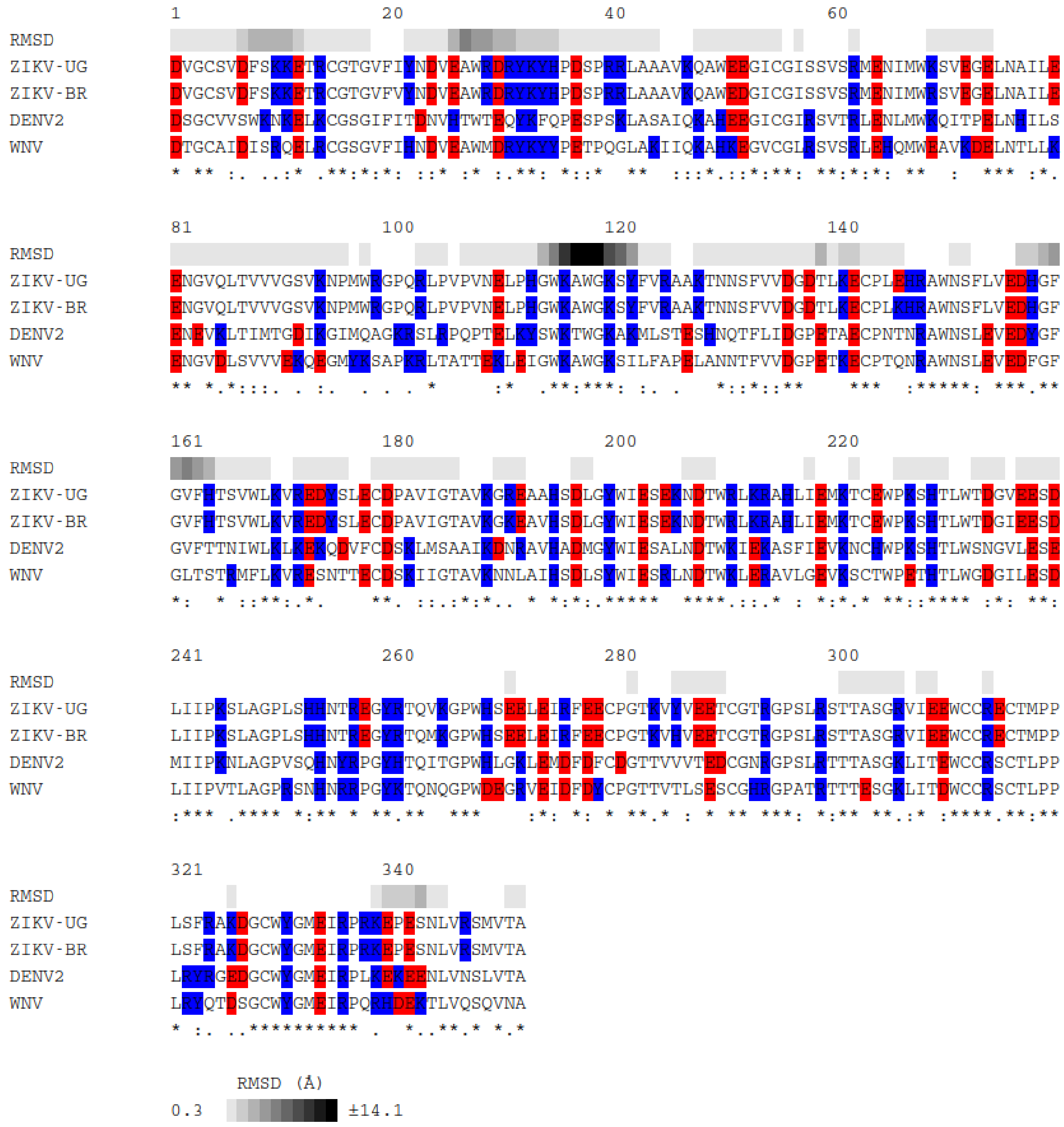


**Figure S1.** Multiple sequence alignment for NS1 proteins. Data for ZIKV-UG, ZIKV-BR, DENV2 and WNV. The acidic and basic residues are highlighted in red and blue, respectively. The first line displays the RMSD among the $NS1_{flavivirus}$ based on their $C_{\alpha}$s per residue. The last line indicates the degree of conservation between proteins. Figure produced with the package *CPRISMA* (Poveda-Cuevas, 2021).

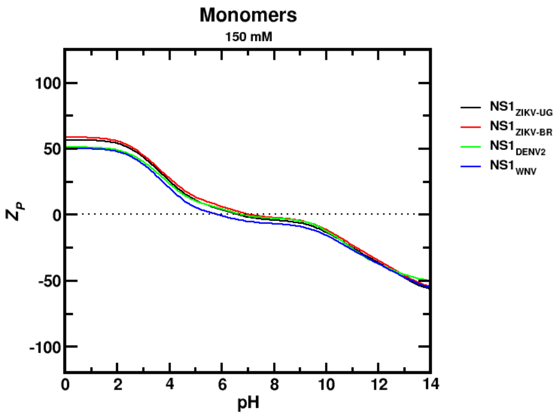
Monomers
150 mM
Z_P
pH
NS1_ZIKV-UG
NS1_ZIKV-BR
NS1_DENV2
NS1_WNV

**(a)**

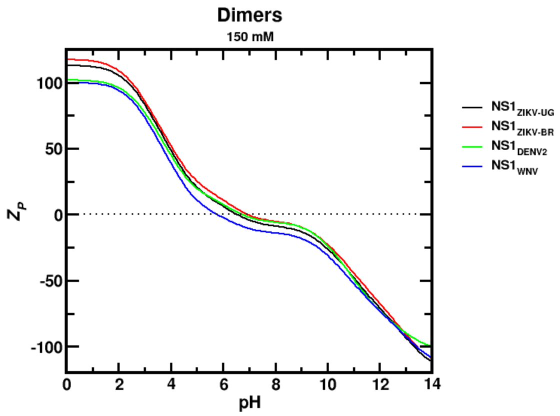
Dimers
150 mM
Z_P
pH
NS1_ZIKV-UG
NS1_ZIKV-BR
NS1_DENV2
NS1_WNV

**(b)**

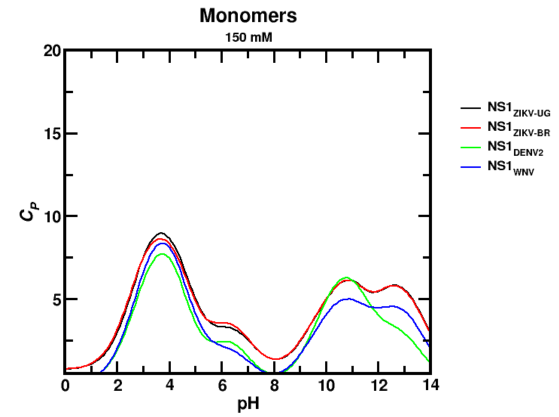
Monomers
150 mM
C_P
pH
NS1_ZIKV-UG
NS1_ZIKV-BR
NS1_DENV2
NS1_WNV

**(c)**

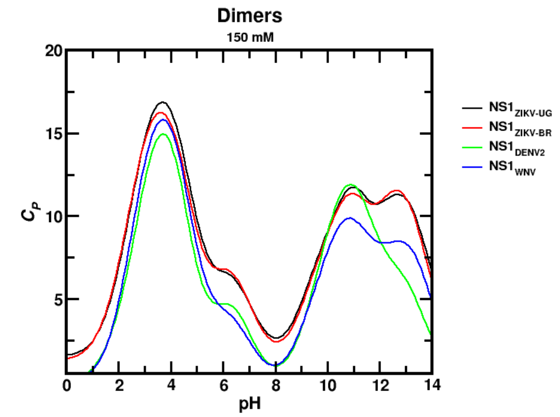
Dimers
150 mM
C_P
pH
NS1_ZIKV-UG
NS1_ZIKV-BR
NS1_DENV2
NS1_WNV

**(d)**

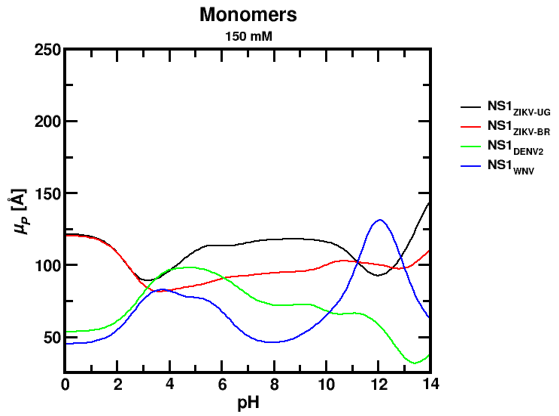
Monomers
150 mM
μ_P [Å]
pH
NS1_ZIKV-UG
NS1_ZIKV-BR
NS1_DENV2
NS1_WNV

**(e)**

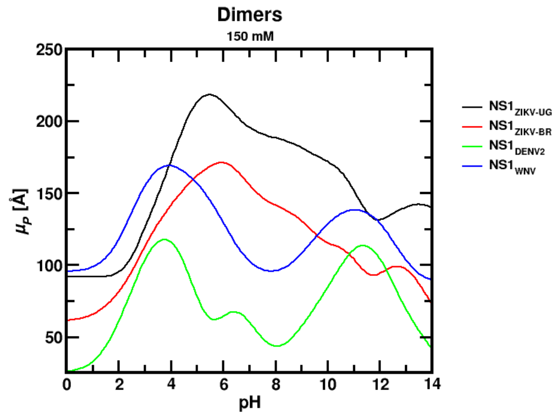
Dimers
150 mM
μ_P [Å]
pH
NS1_ZIKV-UG
NS1_ZIKV-BR
NS1_DENV2
NS1_WNV

**(f)**

**Figure S2.** Main electrostatic features of the NS1 proteins as a function of pH. Data for $NS1_{ZIKV-UG}$ (PDB id 5K6K), $NS1_{ZIKV-BR}$ (PDB id 5GS6), $NS1_{DENV2}$ (PDB id 4O6B), and $NS1_{WNV}$ (PDB id 4O6D) at two oligomeric states. (a,b) Protein net charge number $Z_P$. (c,d) charge regulation capacity ($C_P$). (e,f) Dipole moment number ($\mu_P$). Data from CpH MC simulations at 150 mM of salt concentration. These calculations correspond to sets A (monomers) and B (dimers) described in the Methodology section.

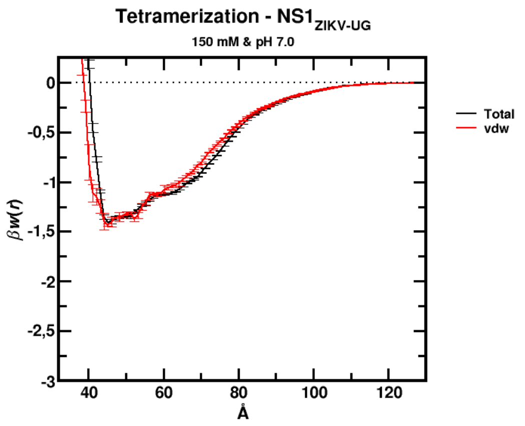


**(a)**

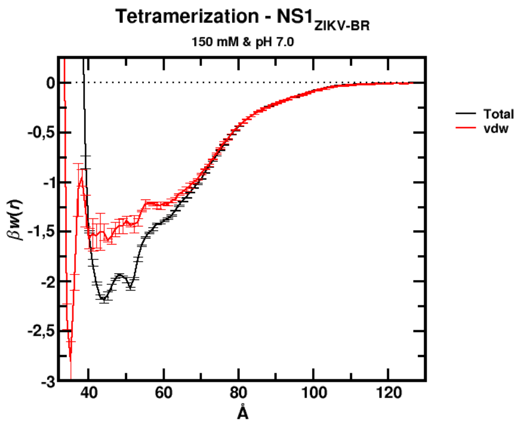


**(b)**

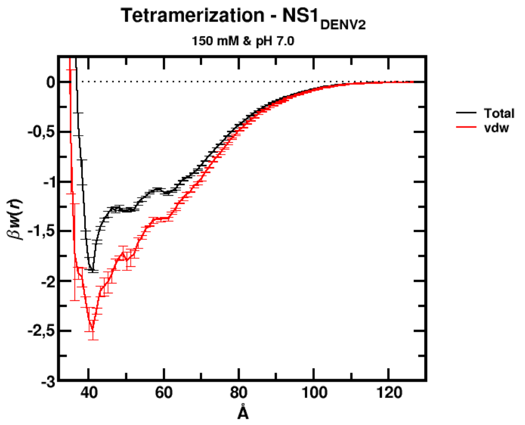


**(c)**

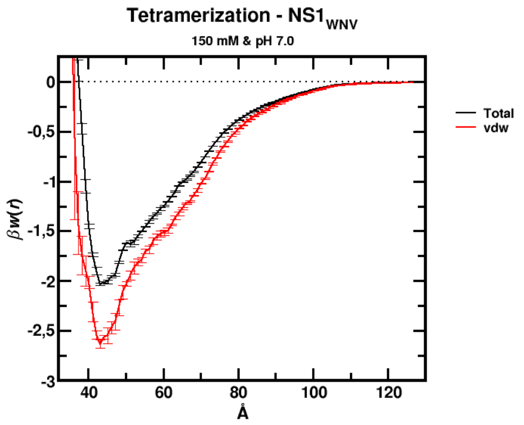


**(d)**

**Figure S3.** Free energy of interactions as a function of the center-center separation distance for the tetramerization of four flavivirus NS1 proteins. The red lines are from an artificial neutral model used solely to evaluate the vdw contributions in the absence of electrostatic interactions. The solid curves in black (total contributions) are the same ones given in Figure 3d where all interactions were incorporated in the model. All other details as in Figure 3. These data correspond to the results from simulation set F.

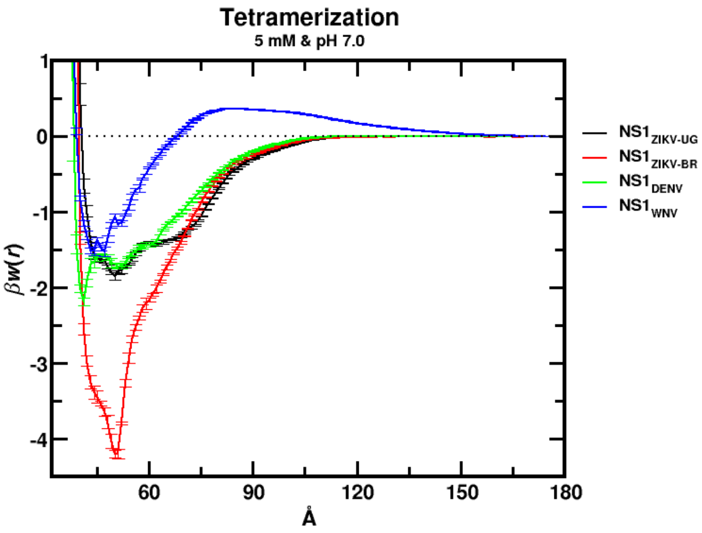


**Figure S4**. Free energy of interactions as a function of the center-center separation distance for the tetramerization of four flavivirus NS1 proteins at low ionic strength. Data from calculations at pH 7 and 5 mM of NaCl. All other details as in Figure 3. These data correspond to the results from simulation set E.
